# The transsulfuration pathway suppresses the embryonic lethal phenotype of glutathione reductase mutants in *Caenorhabditis elegans*

**DOI:** 10.1101/2025.01.09.632117

**Authors:** Marina Valenzuela-Villatoro, David Guerrero-Gómez, Eva Gómez-Orte, Qing Cheng, Angelina Zheleva, José Antonio Mora-Lorca, Dunja Petrovic, Nigel J. O’Neil, Julián Cerón, Akiko Hatakeyama, Shuichi Onami, Milos Filipovic, Elias S.J. Arnér, Juan Cabello, Antonio Miranda-Vizuete

## Abstract

The *gsr-1* gene encodes the only glutathione reductase in *Caenorhabditis elegans* and *gsr-1* loss of function alleles have a fully penetrant embryonic lethal phenotype. Therefore, maintenance of glutathione redox homeostasis is essential for nematode survival. We report here that impairment of the nonsense-mediated mRNA decay (NMD) pathway suppresses the embryonic lethality of *gsr-1* mutants, allowing their normal development and growth. This NMD pathway dependent suppression requires *cth-1* and *cth-2* that encode, respectively, two isoforms of cystathionine-γ-lyase that catalyze the conversion of cystathionine to cysteine through the transsulfuration pathway. Interestingly, the thioredoxin system that can also provide cysteine through the cystine reduction pathway is not required for the suppression of the lethal phenotype of *gsr-1* embryos when the NMD pathway is inactivated. Together, our data indicate that increasing the activity of the reverse transsulfuration pathway can compensate the detrimental effect of the *gsr-1* mutation, raising the interesting question of why *C. elegans* has not preserved such compensatory mechanism to avoid the embryonic lethality of these mutants.

**Highlights:**

- A novel cryptic deletion in the *smg-3* gene suppresses the embryonic lethality of *C. elegans gsr-1* mutants.
- Inactivation of the nonsense-mediated mRNA decay (NMD) pathway allows survival of *gsr-1* mutant worms.
- Genetic impairment of the transsulfuration pathway restores the embryonic lethal phenotype in *gsr-1; smg-3* mutants.

## Introduction

The tripeptide glutathione (GSH: L-γ-glutamyl-L-cysteinyl-glycine) is the most abundant low molecular weight thiol in the vast majority of organisms and it plays key roles in many cellular processes including cell signaling, DNA synthesis and repair, regulation of protein function, detoxification or antioxidant defense. GSH performs its diverse functions either directly through non-enzymatic reactions or as cosubstrate in enzyme-catalyzed reactions (1). Glutathione is synthesized in the cytoplasm of eukaryotic cells by the sequential action of two ATP requiring enzymes: gamma-glutamylcysteine synthetase that catalyzes the condensation of glutamate and cysteine, the rate limiting reaction of GSH biosynthesis, and glutathione synthetase which catalyzes the conjugation of gamma-glutamylcysteine with glycine (2). The relevance of GSH as an essential metabolite is illustrated by the fact that mutations that impair γ-glutamylcysteine synthetase function cause lethality in all organisms studied from yeast to mammals (3–7). Similarly, genetic inactivation of glutathione synthetase is also lethal in *Caenorhabditis elegans* and mice (8,9). On the other hand, no overt phenotype in yeast under normal growth conditions is observed, probably due to the increased levels of gamma-glutamylcysteine that may partially compensate the GSH requirement for growth (10). In humans, the very few cases of patients with glutathione synthetase deficiency have been found to have increased levels of γ-glutamylcysteine, supporting this compensatory role on GSH requirement (11,12).

Once GSH serves as an electron donor it becomes oxidized (GSSG) and is then recycled to its reduced form by the flavoenzyme glutathione reductase, which is not needed for yeast, zebrafish or mice survival (13–15). The dispensability of glutathione reductase in these organisms has been explained by the thioredoxin system working as an alternative pathway to regenerate reduced glutathione (13,16,17), although the exact mechanism needs yet to be fully elucidated. In contrast to yeast, zebrafish and mice, glutathione reductase is essential for *Caenorhabditis elegans* development: homozygous *gsr-1* animals segregating from heterozygous parents (hereafter referred as *gsr-1(m+,z-)*; *m*, maternal and *z*, zygotic) have a normal embryonic and postembryonic development and reach adulthood indistinguishably of wild type controls thanks to the maternally contributed *gsr-1* mRNA and/or GSR-1 protein. In turn, the *gsr-1(m-,z-)* embryos generated by these *gsr-1(m+,z-)* adults arrest at the pregastrula stage due to lack of maternal contribution (8). This embryonic lethal phenotype is intriguing as it has been shown that cytoplasmic thioredoxin reductase TRXR-1 and glutathione reductase GSR-1 have redundant functions in the nematode molting cycle, probably by TRXR-1 mediating a thioredoxin-dependent reduction of GSSG in the hypodermis in the absence of GSR-1 (18). Why TRXR-1 can functionally substitute GSR-1 to allow molting of *gsr-1(m+,z-)* larvae but not to allow *gsr-1(m-,z-)* embryos development is currently unknown.

When GSSG reduction is compromised in mammalian cells by simultaneous blockage of glutathione reductase and cytoplasmic thioredoxin reductase 1, cells rely on *de novo* glutathione synthesis for survival using the reverse transsulfuration pathway to provide cysteine as a GSH precursor (19). This pathway employs homocysteine (generated from methionine by the S-adenosylmethionine cycle) to synthesize cysteine in two sequential enzymatic steps catalyzed by cystathionine β-synthase and cystathionine γ-lyase, respectively (20). Although mammals have a second independent pathway able to supply cysteine from extracellular cystine (oxidized form of cysteine) in a reaction catalyzed by the cystine reductase TRP14 (21), this pathway is not operative in *TRXR1* knockout cells, because TRP14 is a member of the thioredoxin family that requires TRXR1 and NADPH to be maintained in its reduced active conformation (22). Of note, while the transsulfuration and cystine reduction pathways have been relatively well characterized in mammals (20,23), very little is known on their functional roles in other model organisms like *Drosophila melanogaster* or *Caenorhabditis elegans*.

The nonsense-mediated mRNA decay pathway (NMD) is a surveillance mRNA system that was initially discovered as a mechanism to degrade transcripts that contained premature stop codons but was later found to eliminate aberrant mRNAs that cannot be translated by other different reasons, like unproductively spliced mRNAs or mRNAs with very long 3’-untranslated regions (reviewed in (24)). Importantly, whole transcriptomic analyses have widened this view by showing that NMD regulates the dynamics and stability of a large proportion of the eukaryotic transcriptome in all organisms (25–28). The core components of the NMD pathway were first identified in yeast and named UPFs (up frameshift proteins) (29), but have been shown to be conserved across metazoan evolution: UPF1 is an ATP-dependent RNA helicase; UPF2 acts as a molecular bridge between UPF1 and UPF3, promoting UPF1 RNA unwinding and helicase activity while UPF3 binds to exon junction complexes (30,31). In *C. elegans*, the molecular constituents of the NMD pathway were originally discovered as a class of extragenic suppressor genes that, when mutated, share a common phenotype of abnormal morphogenesis of the male bursa and the hermaphrodite vulva, thus named *smg* (**<u>s</u>**uppressor with **<u>m</u>**orphogenetic effect on **<u>g</u>**enitalia) genes (32). This gene class was later found to encode the orthologues of yeast UPF genes: SMG-2/UPF1, SMG-3/UPF2 and SMG-4/UPF3 (33–36). In addition to these core components, there are other proteins that assist for the correct function and regulation of the NMD pathway in all organisms (31,37,38).

In this work, we report that impairment of the NMD pathway suppresses the embryonic lethal phenotype of *C. elegans gsr-1* loss of function mutants and that this suppression requires the two cystathionine γ-lyase orthologues CTH-1 and CTH-2.

## Results

### Inactivation of the NMD pathway suppresses the embryonic lethality of *C. elegans gsr-1* mutants

The *C. elegans gsr-1(tm3574)* deletion allele encodes truncated versions of mitochondrial (GSR-1a) and cytosolic glutathione reductase (GSR-1b) isoforms that lack part of their respective NADPH and FAD binding domains (Figure 1a,b) and *gsr-1(tm3574)* mutants exhibit a fully penetrant embryonic lethal phenotype (Figure 1c) (8). When generating a strain combining the *gsr-1(tm3574)* allele with the integrated transgene *gnaIs2 [Pmyo-2::yfp; Punc-119::Aβ1-42]* (that expresses human Aβ peptide in all worm neurons along with YFP in pharynx as fluorescence co-injection marker) (39), we were able to isolate double *gsr-1(tm3574); gnaIs2* homozygous animals that were viable and grossly wild type (Figure 1c). As the *gnaIs2* transgene in heterozygosis failed to rescue the *gsr-1* embryonic lethal phenotype (Figure 1c), we concluded that the causative suppressor mutation is recessive. The rescue of the *gsr-1(tm3574)* embryonic lethal phenotype by the *gnaIs2* transgene is not due to YFP or Aβ as the control transgenes *gnaIs1 [Pmyo-2::yfp],* expressing YFP in pharynx, or *dvIs50 [Psnb-1:: Aβ1-42],* expressing Aβ in neurons, were unable to restore viability of *gsr-1(tm3574)* mutants (Figure 1c). Ten times outcrossing of the *gnaIs2* transgene with the wild type strain maintained the viability of *gsr-1(tm3574*) mutants, suggesting that the transgene insertion locus or a closely linked mutation was responsible for the suppression of the embryonic lethal phenotype.

**Figure 1.**
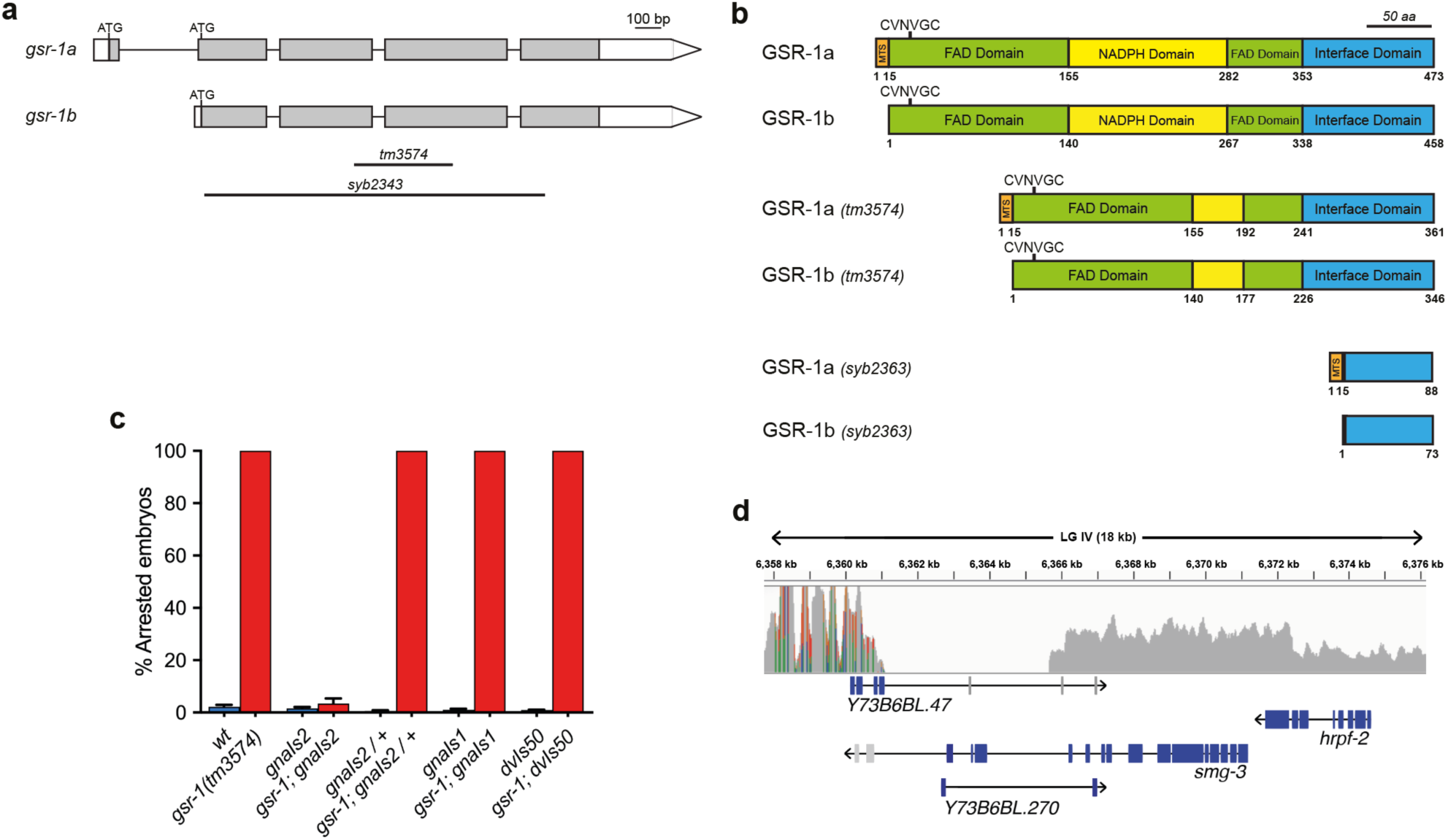
mRNA and protein domain organization of wild type and mutant *gsr-1* and identification of the molecular lesion segregating with the *gnaIs2* transgene. **a) Schematic representation of the two *gsr-1* mRNA variants.** Boxes represent exons and lines show spliced introns. White boxes indicate 5’-UTR and 3’-UTR, respectively, and grey boxes indicate the ORF. Boundaries of *gsr-1(tm3574)* and *gsr-1(syb2343)* deletions are shown as black lines. **b) Schematic representation of GSR-1 proteins**. Protein domain organization of GSR-1a and GSR-1b isoforms as well as those of the shorter proteins resulting from translation of the *gsr-1(tm3574)* and *gsr-1(syb2343)* deletion alleles**. c) The *gnaIs2* transgene suppresses *gsr-1* mutants embryonic lethality.** *gsr-1(tm3574)* embryos carrying the *gnaIs2* transgene (expressing YFP in pharynx and Aβ in neurons) in homozygosis develop normally. The *gnaIs2* transgene in heterozygosis and control transgenes *gnaIs1* (expressing YFP in pharynx) and *dvIs50* (expressing Aβ in neurons) do not suppress *gsr-1* mutants embryonic lethality. Data are the mean +/− SEM of three independent experiments (three biological replicates) with at least 100 embryos laid per plate. Strains with the *wild type gsr-1* allele are depicted in blue and strains with the *gsr-1* mutant allele are depicted in red. **d) Diagram of the deletion boundaries at 6.36 MB position of LG IV identified in the strain harboring the *gnaIs2* integrated transgene.** Genes within this region are represented in boxes where ORF exons are in blue and 3’-UTR exons are in grey, connected by lines representing the introns.

Next, to identify the suppressor locus, we performed whole genome sequencing using a single-nucleotide polymorphism (WGS-SNP) mapping strategy (40). Briefly, the *gnaIs2* harbouring strain was crossed with the highly polymorphic Hawaiian *C. elegans* strain CB4856 and the resulting F2 progeny carrying the *gnaIs2* transgene in homozygosis was pooled for WGS-SNP analysis. The loss of Hawaiian SNPs identified the chromosomal interval containing the suppressor locus within a wide region of LG IV (Supplemental Figure 1). A detailed inspection of this interval identified a 4.5 kb deletion, centered at 6,364 MB position of LG IV (Figure 1d), which harbors three genes: *Y73B6BL.47* and *Y73B6BL.270* of unknown function and *smg-3* that encodes the *C. elegans* orthologue of human UPF2, a core regulator of the non-sense mRNA mediated decay (NMD) pathway (34).

*C. elegans* hermaphrodites with mutations in *smg* genes display a **<u>p</u>**rotruding **<u>vu</u>**lva (Pvu) phenotype (32). Interestingly, the original strain carrying the *gnaIs2* transgene as well as the viable *gsr-1(tm3574); gnaIs2* animals exhibit this Pvu phenotype, similar to that of *smg-3(ma117)* mutants used as control (Figure 2a), suggesting that animals bearing the *gnaIs2* transgene may have impaired *smg-3* function. To determine whether defective SMG-3 function allows *gsr-1* embryos development, we generated double mutants of the *gsr-1(tm3574)* deletion with three independent *smg-3* alleles: *ma117* whose molecular lesion is unknown and the deletions *tm5719* and *tm5906*. We found that all *smg-3* alleles rescued the *gsr-1* embryonic lethal phenotype (Figure 2b).

**Figure 2.**
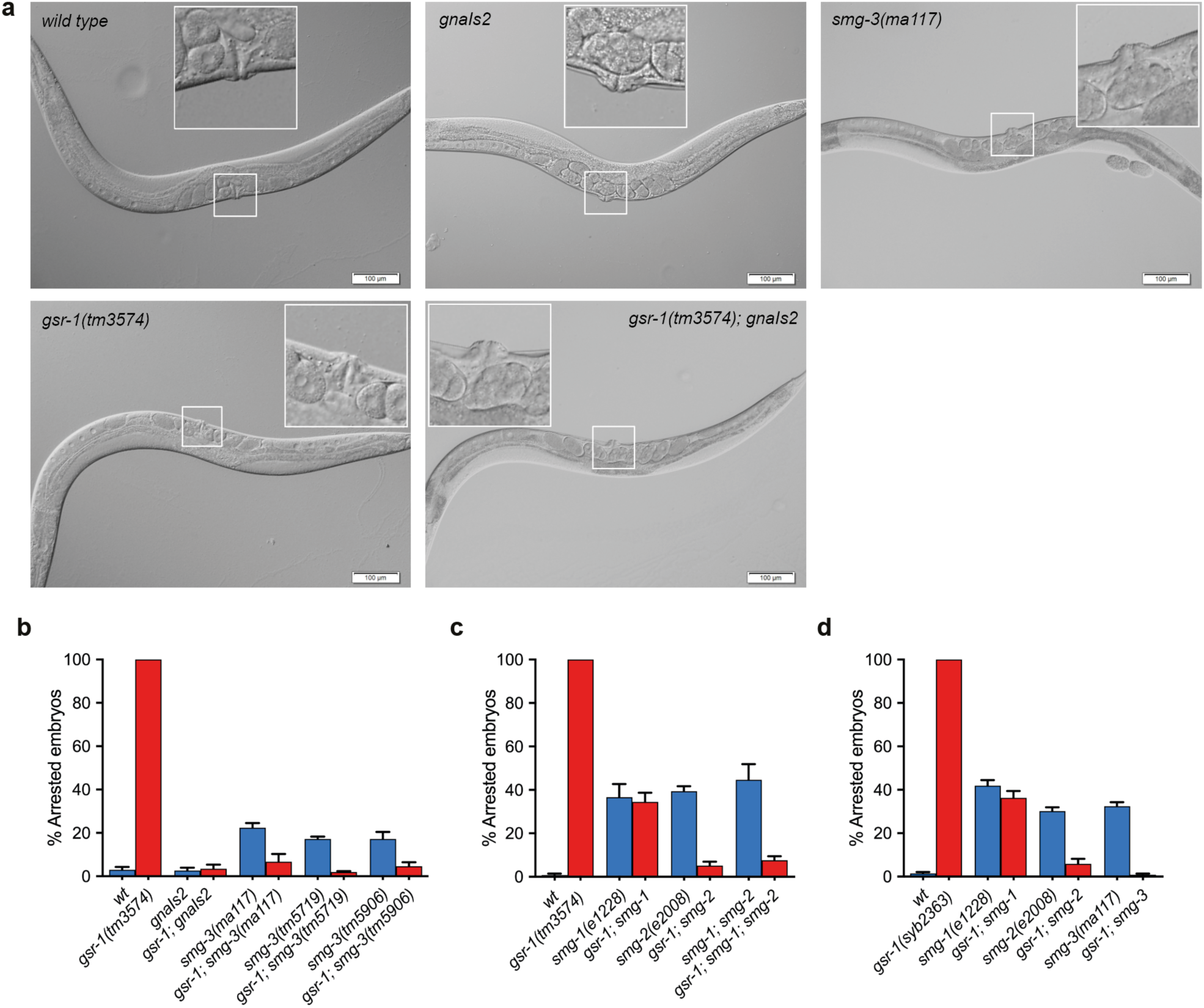
Impairment of the NMD pathway suppresses the embryonic lethal phenotype of *gsr-1* mutants. **a) Representative micrographs of first day adult *gsr-1(tm3574), gnaIs2* or *smg-3(ma117)* mutants.** All mutants develop similar to wild type control, except that those carrying the *gnaIs2* transgene or smg-3 mutation have a protruding vulva phenotype (Pvu). Insets show the magnification of the vulva area**. b,c,d) *smg-1(e1228)*, *smg-2(e2008)* and *smg-3(ma117)* mutations allow normal embryonic development of *gsr-1(tm3574)* and *gsr-1(syb2363)* animals.** The variable number of arrested embryos in the strains carrying the *smg-1*, *smg-2* and *smg-3* alleles is due to the *him-2(e1065)* or *him-5(e1490)* (high incidence of males) mutations present in their respective backgrounds (41) **(See Supplemental Table 1 for complete genotype).** In all cases, the number of arrested embryos substantially decreased upon crossing with the *gsr-1* mutation, including the *smg-1(e1228)* allele that is linked to the *him-2(e1065)* mutation at LG I. Data are the mean +/− SEM of three independent experiments (three biological replicates) with at least 100 embryos laid per plate. Strains with the *wild type gsr-1* allele are depicted in blue and strains with the *gsr-1* mutant allele are depicted in red.

To confirm that impairment of NMD pathway activity is the underlying cause of the *gsr-1* embryonic lethal phenotype suppression, we combined the *gsr-1(tm3574)* deletion with mutations in *smg-1* and *smg-2* genes, that encode other components of the NMD pathway acting upstream *smg-3* (34–36). As shown in Figure 2c, both *smg-1(e1228)* and *smg-2(e2008)* mutants also restored the viability of *gsr-1* mutants. As described above, the *gsr-1(tm3574)* deletion allele encodes shorter GSR-1 isoforms (both cytoplasmic and mitochondrial) that lack part of the NADPH and FAD binding domains (Figure 1a,b) and we have shown that this shorter isoforms are devoid of glutathione reductase enzymatic activity *in vitro* (8). Although the *smg* genes were initially defined as a class of extragenic suppressors (32), given the role of NMD in the regulation of the dynamics and stability of many transcripts (25–28), we first set to rule out that the suppression phenotype would arise from a cryptic expression of the *gsr-1(tm3574)* transcript, producing a shorter GSR-1 protein with residual enzymatic activity *in vivo*. For this purpose, we generated by CRISPR-Cas9 a new deletion allele, *gsr-1(syb2363)* that almost completely eliminates the *gsr-1* ORF, thus being a putative null allele (Figure 1a,b), Similar to *gsr-1(tm3574)* allele, homozygous *gsr-1(syb2363)* animals also display a fully penetrant embryonic lethal phenotype which is suppressed by mutations in the genes encoding the different components of *C. elegans* NMD pathway (Figure 2d), indicating that the suppressor phenotype is extragenic to *gsr-1*. Collectively, these data demonstrate that impairment of the NMD pathway function suppresses *gsr-1* mutants embryonic lethality. This is most likely achieved by allowing the translation of one or more mRNAs (normally degraded by a functional NMD pathway) encoding components of an alternative system capable of generating enough GSH to sustain *gsr-1* embryos development.

### The inactivation of the NMD pathway does not suppress the synthetic molting phenotype of *gsr-1(m+,z-); trxr-1* double mutants

Aiming to identify the alternative system that allows the development of *gsr-1* embryos in an *smg-3* (NMD deficient) background, we first focused on the cytoplasmic thioredoxin reductase TRXR-1, that we have previously shown to function redundantly with GSR-1 in the worm molting cycle (18). For this purpose, we used three different *trxr-1* alleles: *trxr-1(sv47)* is a deletion that removes most of *trxr-1* ORF and is probably a null allele, whereas *trxr-1(cer34[Sec666Cys])* and *trxr-1(cer55[Sec666*])* are two different point mutation alleles that eliminate the selenocysteine residue of TRXR-1, required for its enzymatic activity (18,42). Because *smg-3* maps at LG IV, same as *trxr-1*, we decided to investigate the role of *trxr-1* in *gsr-1* mutants development when the NMD pathway is impaired using instead the *smg-2(e2008)* allele, which is equally functional in inactivating the pathway (Figure 2c). In all cases, the *smg-2* mutation did not suppress the synthetic molting defect of *gsr-1(m+,z-); trxr-1* double mutants (Figure 3). Importantly, the very few *gsr-1(m+,z-); trxr-1; smg-2* survivors that managed to overcome the four larval molts became adults with a severe sick appearance that laid very few embryos which invariably arrested at the pregastrula stage, like *gsr-1(m-,z-)* mutant embryos (data not shown) (8). Thus, these data suggest that the functional redundancy between GSR-1 and TRXR-1 in the worm molting cycle is independent of the NMD pathway.

**Figure 3.**
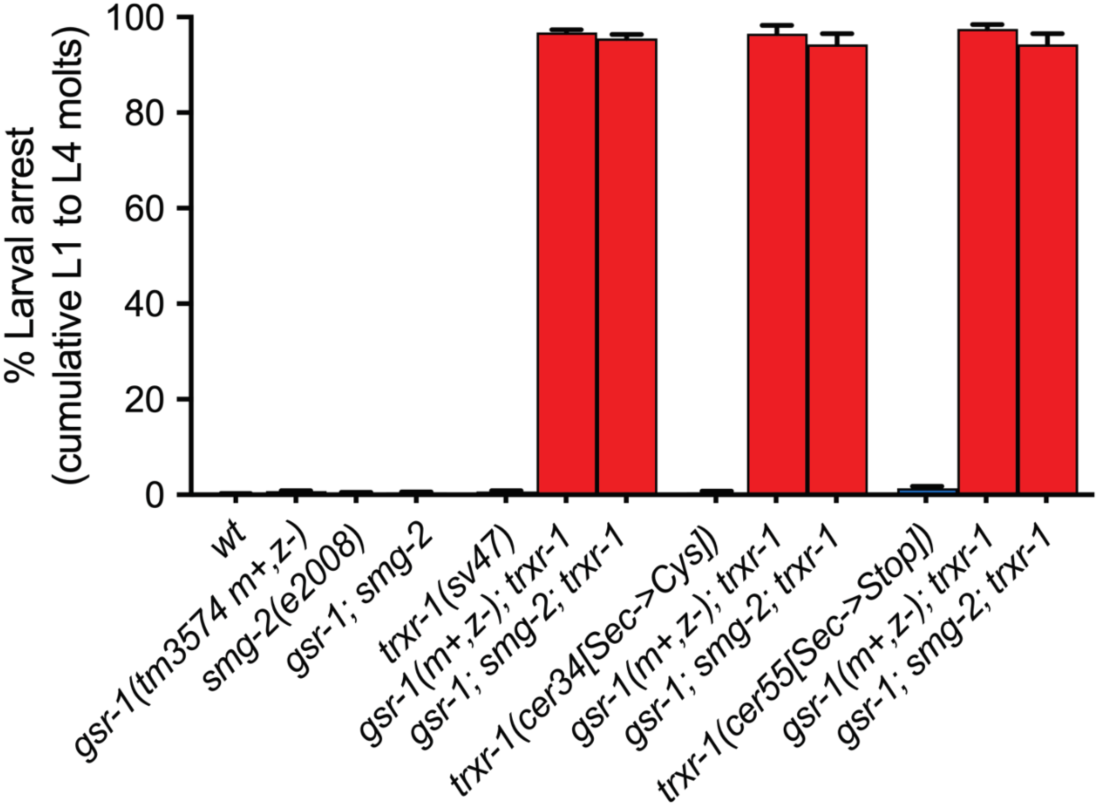
The *smg-2(e2008)* mutation fails to suppress the molting phenotype of *gsr-1; trxr-1* double mutants. All triple mutants *gsr-1; smg-2; trxr-1* display a highly penetrant molting larval arrest phenotype, similar to that of the control strain *gsr-1; trxr-1*. Data are the mean +/− SEM of three independent experiments (three biological replicates) with at least 100 embryos laid per plate. Strains with the *wild type gsr-1* allele are depicted in blue and strains with the *gsr-1* mutant allele are depicted in red.

### The thioredoxin system is dispensable for the growth of *gsr-1; smg-3* embryos

As mentioned above, the *gsr-1(m+,z-); trxr-1; smg-2* triple mutants that make it through the four molting steps are sick and the very few *gsr-1(m-,z-); trxr-1; smg-2* embryos they produce do not develop. To elucidate whether the embryonic arrest of this triple mutant is due to a direct need of the thioredoxin system in reducing GSSG in the absence of GSR-1 or, instead, it is due to other requirements of the worm thioredoxin system during embryogenesis, not directly related to glutathione metabolism, we first aimed to determine whether *C. elegans* TRXR-1 exhibits glutathione reductase enzymatic activity *in vitro*. Despite metazoan thioredoxin reductases and glutathione reductases share domain organization and functional groups, apart from the additional C-terminal active site motif in thioredoxin reductases mentioned above (43), no metazoan thioredoxin reductase has been shown to reduce GSSG *in vitro*. Consistently, this is also the case for *C. elegans* TRXR-1, which we found unable to reduce GSSG in the presence of NADPH (Figure 4a and Supplemental Figure 2). Having ruled out direct GSSG reduction by TRXR-1, we next addressed whether TRXR-1 role in allowing *gsr-1; smg-3* embryos development could be mediated through reduction of one of the nematode thioredoxin family members that have previously been shown to interact with the glutathione system in other organisms. We first focused on the thioredoxin member DPY-11 because is expressed in the hypodermis (where GSR-1 and TRXR-1 display redundant functions in the molting cycle) (44) and its mammalian orthologue TMX1 has been reported to use glutathione for its catalytic activity (45). However, two different putative *dpy-11* null alleles *(e207* and *sy748)* (44,46) and a newly generated CRISPR-Cas9 allele *(syb4162[C51S;C54S])* that encodes a redox-dead DPY-11 variant, all failed to restore embryonic lethality in *gsr-1; smg-3* mutants (Figure 4b). To our surprise, the redox-dead allele *dpy-11(syb4162)* does not cause any dumpy phenotype (Supplemental Figure 3), indicating that DPY-11 enzymatic redox activity is dispensable for its biological function in maintaining body morphology, in contrast to what has been previously proposed (44). Like *dpy-11* mutants, null or loss of function alleles of *trx-1*, *txdc-17* and *txl-1* genes, whose mammalian orthologs encode the thioredoxin proteins with GSSG reducing activity *in vitro* TRX-1 (47), TRP14/TXNDC17 (21) and TXNL1/TRP32 (48) had not effect on the development of *gsr-1; smg-3* mutants (Figure 4c). Together, these data suggest that the thioredoxin system is not directly involved in the mechanism by which impairment of the NMD pathway suppresses the embryonic lethality of *gsr-1* mutants. However, given the high number of thioredoxin family members in *C. elegans* (49) and their possible functional redundancy in redox mediated reactions, our data cannot completely rule out that one or more yet unidentified TRXR-1 dependent thioredoxins may play a role in the viability of *gsr-1; smg-3* animals.

**Figure 4.**
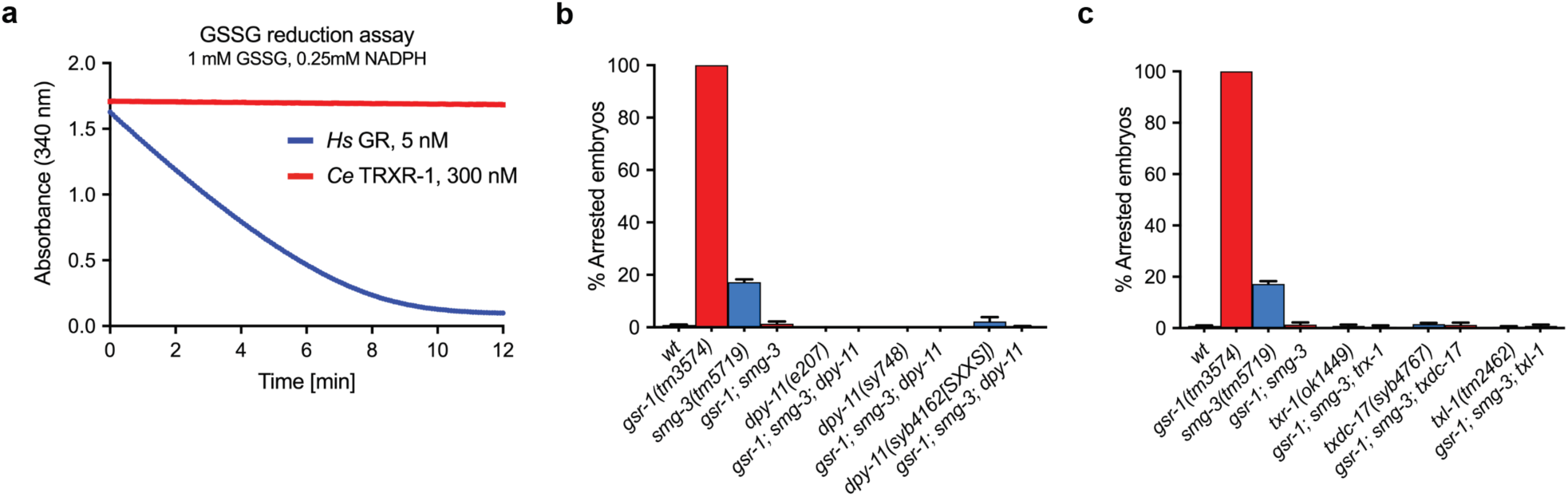
The thioredoxin system is not required for the suppression of *gsr-1* embryonic lethality upon impairment of the NMD pathway. **a) *C. elegans* TRXR-1 is devoid of GSSG reducing activity *in vitro*.** NADPH consumption in the presence of 1 mM GSSG of recombinantly expressed human glutathione reductase (*Hs* GR) and *C. elegans* thioredoxin reductase 1 (*Ce* TRXR-1). **b, c) The thioredoxins DPY-11, TRX-1, TXDC-17 and TXL-1 are not required for the embryonic development of *gsr-1; smg-3* animals.** All triple mutants combinations of genes encoding thioredoxin proteins with GSSG reducing activity in a *gsr-1; smg-3* doble mutant background develop similarly as *gsr-1; smg-3* control animals Data are the mean +/− SEM of three independent experiments (three biological replicates) with at least 100 embryos laid per plate. Strains with the *wild type gsr-1* allele are depicted in blue and strains with the *gsr-1* mutant allele are depicted in red.

### The transsulfuration pathway supports development of *gsr-1; smg-3* double mutants

We have previously shown that *C. elegans* expressing aggregation-prone proteins in muscle cells are extremely sensitive to GSH depletion (50) and that these animals rely on functional transsulfuration and cystine reduction pathways for survival, most likely by provisioning cysteine for *de novo* GSH synthesis (23). Cystine reduction is performed by TRXR-1 and TRP14/TXDC-17 in both mammals and *C. elegans* (23). However, our data indicate that TRXR-1 and TXDC-17 are not required for viability of *gsr-1; smg-2* or *gsr-1; smg-3* mutants (Figure 3 and 4c). Hence, we asked whether the transsulfuration pathway would suffice to supply the necessary cysteine for *de novo* GSH synthesis to allow development of *gsr-1; smg-3* embryos. To test this hypothesis, we employed animals that carry loss of function mutations in the *cth-1* and *cth-2* genes, encoding two paralogues of the enzyme cystathionine γ-lyase that catalyzes the last step in the transsulfuration pathway, converting cystathionine into cysteine and α-ketobutyrate (20). Similar to *txdc-17* mutants, worms bearing mutations either in the *cth-1* or *cth-2* genes did not impair the development of *gsr-1; smg-3* embryos (Figure 5a). However, *gsr-1; smg-3; cth-1; cth-2* quadruple mutant worms produced about 80% of arrested embryos (Figure 5a), demonstrating a functional redundancy of CTH-1 and CTH-2 in the viability of *gsr-1; smg-3* embryos and suggesting that cystathionine to cysteine conversion through transsulfuration is the main pathway responsible for sustaining development of *gsr-1; smg-3* embryos. Interestingly, a *gsr-1; smg-3; txdc-17; cth-1; cth-2* quintuple mutant increased the number of arrested embryos to 93% as compared with the 80% of the *gsr-1; smg-3; cth-1; cth-2* quadruple mutant (Figure 5a), indicating that some cysteine provisioning through the cystine reduction pathway can occur in the absence of a functional transsulfuration pathway, but that it is not enough on its own to allow *gsr-1* embryos survival.

**Figure 5.**
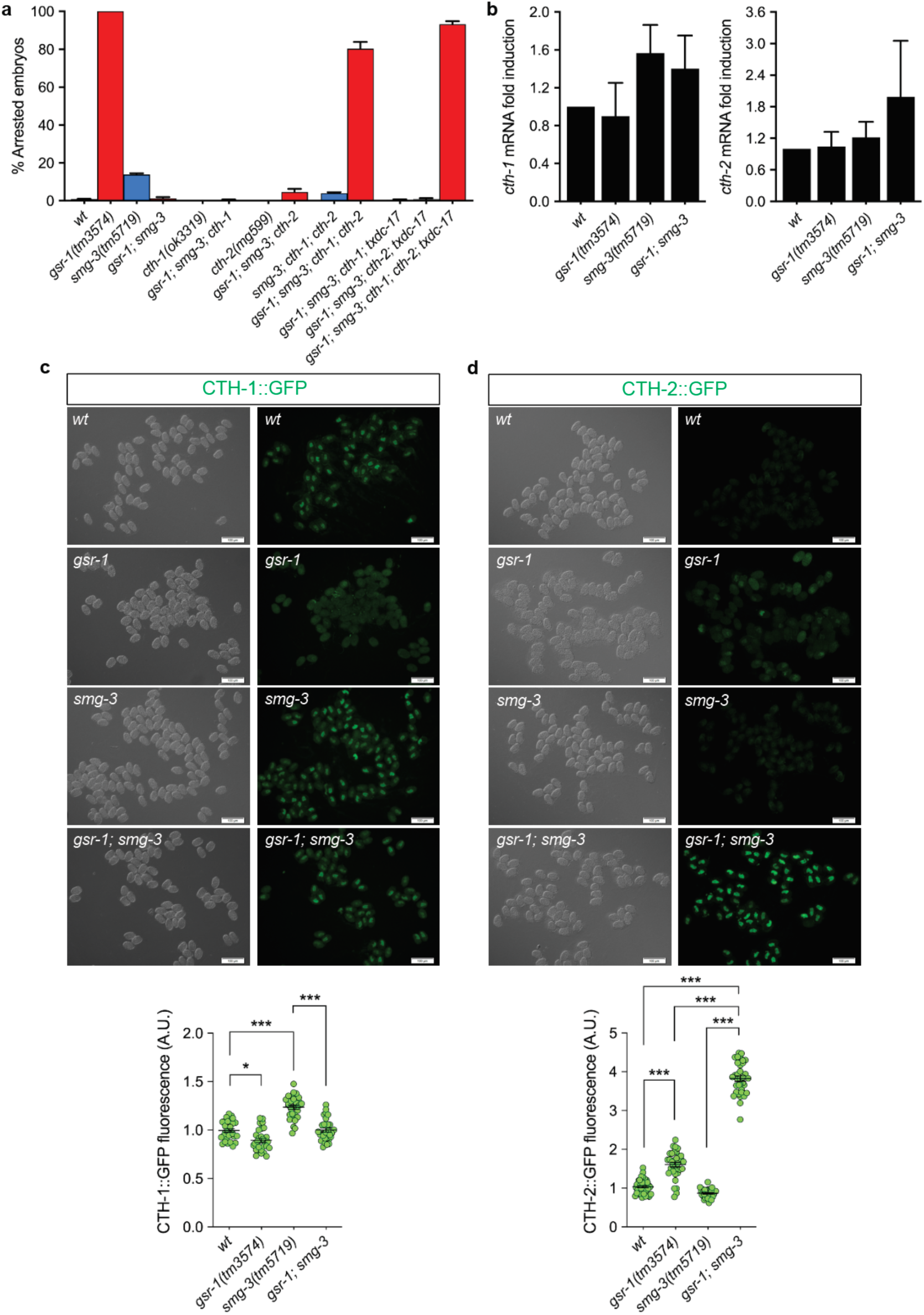
Inactivation of the transsulfuration pathway impairs *gsr-1; smg-3* double mutants development. **a) Simultaneous inactivation of *cth-1* and *cth-2* genes restores the embryonic lethal phenotype in *gsr-1; smg-3* double mutants.** Data are the mean +/− SEM of three independent experiments (three biological replicates) with at least 100 embryos laid per plate. Strains with the *wild type gsr-1* allele are depicted in blue and strains with the *gsr-1* mutant allele are depicted in red**. b) *cth-1* and *cth-2* mRNA levels in *gsr-1* and *smg-3* mutants.** Data are the mean +/− SEM of three independent experiments (three biological replicates). Data were not significantly different by one-way ANOVA multiple comparisons test. **c,d) CTH-1 and CTH-2 protein levels in *gsr-1* and *smg-3* mutants.** Differential interference contrast (left panel) and fluorescence (right panel) microscopy images and quantification of CTH-1::GFP and CTH-2::GFP endogenous reporters in *gsr-1* and *smg-3* mutants. Graphs represent the data of 3 independent experiments with at least 30 embryos. *** p<0.001 by Kruskal-Willis with Dunńs multiple comparisons test. Error bars are SEM.

Because the NMD pathway regulates the dynamics and stability of part of the eukaryotic transcriptome, we reasoned that the *smg-3* mutation may stabilize or increase the levels of *cth-1* and *cth-2* mRNAs, probably resulting in higher levels of protein expression, thus allowing the development of *gsr-1* embryos. To test this hypothesis, we first quantified *cth-1* and *cth-2* transcript levels in embryos of the different genetic backgrounds by qPCR. As shown in Figure 5b, the *smg-3* mutation did not significantly increase the amount of *cth-1* and *cth-2* mRNAs.

Next, to address if the *smg-3* mutation would increase CTH-1 and CTH-2 proteins amount without altering their corresponding mRNA levels, we generated by CRISPR-Cas9 transgenic strains that produce CTH-1::GFP and CTH-2::GFP fusion proteins, expressed from their respective endogenous promoters (Supplemental Table 1). Using these endogenous GFP reporters we found that the *smg-3* embryos only slightly increased the levels of CTH-1::GFP but not those of CTH-2::GFP (Figure 5c and 5d). In contrast, in a *gsr-1; smg-3* double mutant background CTH-1::GFP levels remained similar to those found in *smg-3* single mutants while CTH-2::GFP levels were significantly increased (Figure 5c and 5d). Finally, *gsr-1* mutant embryos slightly decrease CTH-1::GFP while increased CTH-2::GFP levels(Figure 5c and 5d).

Taken together, these data indicate that *gsr-1* embryos do not substantially increase *cth-1* and *cth-2* mRNA and protein levels to compensate for the lack of GSSG reducing capacity. However, impairing NMD pathway activity would result in stabilization of *cth-1* and *cth-2* mRNAs, thus allowing *gsr-1* embryos to bypass the pregastrula stage checkpoint (8) thanks to a sustained provision of CTH-1 and CTH-2.

## Discussion

Glutathione is the most prevalent thiol-based redox metabolite in virtually all organisms, with few exceptions like the low molecular weight thiols bacillithiol or mycothiol that serve similar functions in some Gram-positive bacteria, ergothioneine in some fungi or trypanothionine in euglenozoa (51). Hence, inactivating mutations in genes encoding the first step in glutathione synthesis enzymes are lethal in all those organisms that use glutathione (3–7). In contrast, mutations in the gene encoding glutathione reductase, the enzyme that recycles oxidized glutathione to its reduced form, not always result in lethal phenotypes. For instance, human patients with mutations in the *GSR* gene can reach old age, although they suffer hemolytic anemia, cataracts, deafness and hyperbilirubinemia (52,53). *GSR* knockout mice do not show any phenotype under stabulary conditions (14) but are highly sensitive to bacterial and fungal infections (54,55). Similarly, putative null GSR zebrafish develop similar to wild type control animals (15). The viability of vertebrate *GSR* mutants has been explained by the thioredoxin system acting as a backup pathway to reduce GSSG (16,17), although the embryonic lethality of *TXNRD1* and *TXN1* knockout mice (56,57) has hampered the experimental demonstration with double knockout mice combinations. Consistent with the thioredoxin system acting as an alternative mechanism to maintain GSH pool, *Drosophila melanogaster* and probably other insects do not possess glutathione reductase and GSSG recycling is carried out by dedicated thioredoxin reductase and thioredoxin proteins (58). Also, yeast lacking glutathione reductase *glr1* gene have an absolute requirement of the thioredoxin system for survival (13).

An exception to this rule is the nematode *C. elegans* as *gsr-1* mutants are embryonic lethal (8), despite having a complete thioredoxin system (49). Why *C. elegans gsr-1* mutants are the only eukaryotes that have an embryonic lethal phenotype even when TRXR-1 can functionally substitute GSR-1 in the worm molting cycle (18)? One possibility is that the *trxr-1* gene is not expressed at enough levels to compensate for the lack of *gsr-1* during the first embryonic divisions, as the *gsr-1* embryos arrest at the pregastrula stage (8). This is consistent with the fact that *trxr-1* gene expression gradually increases during the first 200 minutes of embryonic development (59). Alternatively, mammalian cells with strongly impairment of glutathione reductase activity excrete GSSG to the extracellular medium to alleviate its toxic intracellular built-up (60). Should this mechanism also operate in *C. elegans* embryonic cells, the constrain imposed by the embryo eggshell could hamper the survival of the *gsr-1* embryos by avoiding GSSG dilution in the extracellular milieu. To test this hypothesis, we eliminated the eggshell of *wild type* and *gsr-1* embryos by chitinase treatment and found that *gsr-1* embryos lacking eggshell still displayed an embryonic lethal phenotype while *wild type* embryos develop normally. Because *gsr-1* embryos lacking eggshell maintained the integrity of the permeability barrier, GSSG could still accumulate within the peri-embryonic. However, removing the embryo permeability barrier to allow GSSG diffusion provokes *wild type* embryos death, thus precluding any conclusion in this direction (Supplemental Figure 4).

The serendipitous finding of the inactivation of the NMD pathway allowing the development of *gsr-1(m-,z-)* embryos suggested that expression or stabilization of one or more unknown mRNAs as the cause of the suppression of the embryonic lethal phenotype. Given that the thioredoxin system has been shown to act as backup system to reduce GSSG in the absence of glutathione reductase (13,16,17), and that *C. elegans* TRXR-1 cooperates with GSR-1 in the worm molting cycle (18), we first approached to evaluate if the thioredoxin system was behind the suppression of the *gsr-1* embryos lethal phenotype by *smg-1*, *smg-2* or *smg-3* mutations. To our surprise, mutations in the *trxr-1* gene encoding cytoplasmic thioredoxin reductase as well as in *dpy-11*, *trx-1* and *txdc-14* genes, encoding the orthologues of those thioredoxins that have been reported in other organisms to reduce GSSG, failed to restore embryonic arrest in *gsr-1; smg-2* or *gsr-1; smg-3* double mutant backgrounds. Thus, the worm thioredoxin system was not directly involved in the development of *gsr-1* embryos with inactivated NMD pathway function.

Mouse hepatocytes that lack cytoplasmic thioredoxin reductase TRXR1 and glutathione reductase GSR can grow *in vitro* as long as methionine is supplied to allow cysteine production for *de novo* GSH synthesis by the reverse transsulfuration pathway (19). In this pathway, methionine enters the S-adenosylmethionine cycle to generate homocysteine, whose accumulation is toxic in both mammals and worms (61,62). Homocysteine is then converted into cysteine by a two-step enzymatic process: first homocysteine is used by cystathionine β-synthase to generate cystathionine and subsequently, cystathionine γ-lyase converts cystathionine into cysteine and a-α-ketobutyrate (20). While in mammals there is only one enzyme of each class, *C. elegans* has two cystathionine β-synthases CBS-1 and CBS-2 (61,63) and two cystathionine γ-lyases CTH-1 and CTH-2 (64). To evaluate whether the activity of the transsulfuration pathway underlays the viability of *gsr-1; smg-3* embryos we combined this double mutant with *cth-1* and/or *cth-2* mutants and found that only the quadruple mutant *gsr-1; smg-3 cth-1*; *cth-2* restored the embryonic lethality. This result implies that NMD pathway inactivation somehow stabilizes the activity of the reverse transsulfuration pathway to a level that counteracts the absence of GSR-1 to reduce GSSG, probably providing enough cysteine for *de novo* GSH synthesis. The dual requirement of *cth-1* and *cth-2* genes in the survival of *gsr-1* embryos is in sharp contrast with previous data demonstrating non-redundant functions of *C. elegans cth-1* and *cth-2* in molybdenum cofactor transfer from bacteria to worms (64), nematode longevity (65) or muscle proteostasis (23) where only *cth-2* is required. In this last scenario, cystine reduction pathway via TRP14/TXDC-17 and transsulfuration pathway via CTH-2 cooperate to maintain worm viability when muscle proteostasis is compromised (23). However, the viability of *gsr-1; smg-3* embryos is only dependent on the transsulfuration pathway, that requires both CTH-1 and CTH-2. This can be explained by the fact that cystine is mostly incorporated from the extracellular environment. However, given the impermeability of *C. elegans* embryos eggshell, cystine cannot be imported into the embryonic cells, therefore relying exclusively cysteine provision by the transsulfuration pathway.

In summary, we have found that in *C. elegans*, like in mammals, the reverse transsulfuration pathway is able, under certain circumstances, to allow survival of gsr-1(m-,z-) embryos, probably by supplying cysteine for *de novo* GSH synthesis when GSSG reduction is compromised. Why this mechanism is not operative in *gsr-1* mutants is intriguing and may be related to low expression levels or tissue specificity of CTH-1 and/or CTH-2 proteins during the early embryo development, thus causing embryonic lethality by accumulation of the toxic metabolite homocysteine and GSSG.

## Experimental Procedures

### C. elegans strains

The standard methods used for culturing and maintenance of *C. elegans* were as previously described (66). A list of all strains used and generated in this study is provided in Supplemental Table 1. The alleles *gsr-1(syb2363)*, *dpy-11(syb4162), cth-1(syb7115)* and *cth-2(syb7086)* were generated at SunyBiotech (http://www.sunybiotech.com) by CRISPR-Cas9 editing. All VZ strains are 6x outcrossed with N2 wild type, except those strains generated by CRISPR-Cas9 that were 2x outcrossed. Worm reagents and details on the protocols used for genotyping the different alleles reported in this work can be provided upon request.

### Whole genome sequencing single-nucleotide polymorphism (WGS-SNP) mapping

Using the Hawaiian single-nucleotide polymorphism (SNP) mapping method, we backcrossed the GRU102 strain carrying the *gnaIs2* transgene with the polymorphic *C. elegans* Hawaiian strain CB4856 (67). Next, we isolated the newly generated F2 recombinants homozygous for the *gnaIs2* transgene. Total DNA extraction was performed using the Plant/Fungi DNA Isolation Kit (Norgen Biotek Corp). Sequencing libraries were constructed using the NEXTflex Rapid DNA-Seq Kit according to manufactureŕs instructions (Bioo Scientific). DNA quality and integrity were evaluated by Experion Automated Electrophoresis System (Bio-Rad) and the concentration was calculated using qPCR. Libraries were prepared at the Genomic Platform at CIBIR (http://cibir.es/es/plataformas-tecnologicas-y-servicios/genomica-y-bioinformatica) and sequenced on an Illumina HiSeq15000. The quality of DNAseq results was assessed using FastQC (http://www.bioinformatics.babraham.ac.uk/projects/fastqc/). The FastQ files were analyzed using a Cloud-Based Pipeline for Analysis of Mutant Genome Sequences (Cloudmap tool, https://usegalaxy.org/cloudmap) with standard parameters following Cloudmap workflow (68).

### Embryonic arrest phenotype

All experiments were performed on synchronized embryos generated by allowing 10 to 15 gravid hermaphrodites to lay eggs during 2.5 hours on seeded plates at 20°C. After parent removal, laid embryos were counted and plates were further incubated at 20°C. During the next two days arrested embryos are count and the number of arrested embryos at day 2 were used for quantification. Viable progeny was quantified on the plates that were incubated for one or two more days (depending on the strain) at days at 20°C.

### Larval arrest phenotype

Synchronized animals were generated by allowing 10 to 15 gravid hermaphrodites to lay eggs during 2.5 hours on seeded plates at 20°C. After parent removal, laid embryos were counted and further incubated for four days at 20°C. For quantification, all larvae were censored and only animals that passed the L4 molt and became young adults were counted.

### Recombinant expression and enzymatic characterization of *C. elegans* TRXR-1

The amino acid sequence encoding the *C. elegans* thioredoxin reductase 1 (TRXR-1) was retrieved from the GenBank database (Accession No. AAD41826.1). The open reading frame (ORF) was subsequently synthesized by Integrated DNA Technologies (IDT) with codon optimization for enhanced expression in *E. coli*. Additionally, an N-terminal hexahistidine-tagged Small Ubiquitin-like Modifier (H6SUMO) fusion was engineered upstream of TRXR-1, and a selenocysteine insertion sequence (SECIS) element was placed downstream. This construct was cloned into the pABC2 vector as previously described (69), yielding the plasmid pABC2a-*Ce*TRXR-1, which was subsequently transformed into the *E. coli* C321.ΔA strain (70).

For the protein expression, a single bacterial colony was cultured in 10 ml of Terrific Broth (TB) at 30°C with constant shaking at 250 rpm overnight. The resultant culture was then scaled up to 2 liters of TB, and incubation was continued under the same conditions until the optical density at 600 nm (OD600) reached 1-1.5. At this point, 5 µM sodium selenite and 0.5 mM IPTG were added into the culture, which was then incubated at a reduced temperature of 24°C overnight.

The bacterial cells were harvested by centrifugation at 5,000 × g for 15 minutes and resuspended in IMAC binding buffer (50 mM Tris-HCl, 250 mM NaCl, and 20 mM imidazole). The cells were then lysed, and the lysate was centrifuged at 30,000 × g for 30 minutes at 4°C. The resulting supernatant was subjected to affinity chromatography purification. The *Ce*TRXR-1 protein was engineered with an N-terminal His-tagged SUMO tag (H6SUMO). The H6SUMO-*Ce*TRXR-1 was initially purified using IMAC (ÄKTAExplorer 10 FPLC equipped with a 5 ml HisTrap FF column, GE Healthcare). To the eluted fraction containing H6SUMO-*Ce*TRXR-1, 15 µg/ml His-tagged ULP1 (SUMO protease) was added, and the protein mixture was incubated at 4°C overnight with gentle shaking to cleave the *Ce*TRXR-1. Excess imidazole in the digestion mixture was removed using a NAP-25 desalting column (GE Healthcare), and the protein mixture was subjected to a second round of IMAC. During this step, non-cleaved H6SUMO-*Ce*TRXR-1, the cleaved H6SUMO tag, His-tagged ULP1, and any impurities from the first round of IMAC bound to the nickel column, while the non-tagged *Ce*TRXR-1 was collected in the flow-through. The purified protein was concentrated and stored in TE buffer containing 30% glycerol for long-term storage at −20°C. Protein concentration was determined spectrophotometrically by measuring FAD absorption at 463 nm (ε = 13,600 M⁻¹cm⁻¹, with one FAD corresponding to one *Ce*TRXR-1 subunit).

Enzyme activity assays were conducted using previously established protocols (71,72). Specifically, the NADPH-dependent reduction of 5,5’-dithiobis(2-nitrobenzoic) acid (DTNB) was employed to determine the specific activities of *Ce*TRXR-1. This assay involves the reduction of one DTNB molecule to two 5-thio-2-nitrobenzoic acid (TNB) molecules by *Ce*TRXR-1. TNB exhibits a strong absorbance at 412 nm, allowing the reaction velocity to be quantified as µmol DTNB reduced per minute, calculated using the extinction coefficient for TNB (13,600 M⁻¹cm⁻¹). To further verify Sec-dependent activities of *Ce*TRXR-1, the alternative insulin-coupled Trx reduction assay was used. Insulin is a substrate of human Trx1 that can be recycled by *Ce*TRXR-1 in the consumption of NADPH. This assay therefore monitored the consumption of NADPH through the decreased absorbance at 340 nm, using NADPH’s extinction coefficient of 6,200 M⁻¹cm⁻¹. For measuring human glutathione reductase (GR) activity, the assay utilized GSSG as the substrate, which was reduced by GR using NADPH, which can be monitored similarly.

### Microscopy

For protruding vulva phenotype analysis (Fig. 2a), second day adult animals were used. For CTH-1::GFP and CTH-2::GFP fluorescence determination in embryos (Fig. 5c,d) an one hour egg lay was performed and then allow the embryos to develop 2 more hours at 20°C on the plate to match the stage at which *gsr-1* embryos arrest. Next, embryos are transferred to a 3% agarose pad on a microscope slide and imaged. Differential interference contrast and fluorescence imaging was performed in an Olympus BX61 microscope equipped with a DP72 digital camera coupled to CellSens Software for image acquisition and analysis. Photoshop CC 2018 and Adobe Illustrator software were used to produce figures. ImageJ Software was used to quantify the fluorescence of the embryos.

### qPCR analysis

To determine the transcriptional activity of *cth-1* and *cth-2* genes in embryos, young adult worms were collected and bleached, to use only embryos at the early stage of division. This way they can be compared with *gsr-1* embryos that arrest between 15 and 120 cells stage (8). For total RNA extraction, eggs were lysed using mirVana Kit, following manufactureŕs instructions (Ambion). Homogenization of the lysate was performed using a conventional rotor-stator homogenizer polytron, pre-chilled with liquid nitrogen. For cDNA synthesis, RNA was extracted with DNAse to eliminate any DNA contamination. In each sample, a total reaction of 10 µl contained: 500ng RNA, 1 µl RQ1 RNase-Free DNAse (Promega), 1x RQ1 DNAse 10x Reaction Buffer and DEPC water. Reactions were incubated for 30 min at 37°C and then stopped by adding 1 µl STOP solution (Promega) and further incubation for 15 min at 65°C. cDNA synthesis was performed using SuperScript III First-Strand Synthesis System for RT-PCR (Invitrogen) following instructions for random hexamers primed. cDNA was eluted in TE buffer.

For qRT-PCR analysis Power SYBR Green Master Mix (ThermoFisher Scientific) and specific primers were used in a QuantStudio 5 Real Time PCR System (Applied Biosystems, ThermoFisher). Normalization to actin *act-1* expression was used to calculate relative expression. The experiments were carried out in three independent replicas. Primers pairs used were (5’->3-): *cth-1*Fw: ATGACTCCGTACTTCCAGCG; *cth-1*Rv: TAGATGCTGCTGGAACTCGT; *cth-2*Fw: GCCGTGTTGCTGTTCCTAAT; *cth-2*Rv: CCACTGCGGCAATATCAACA; *act-1*Fw: ACGCCAACACTGTTCTTTCC; *act-1*Rv: GATGATCTTGATCTTCATGGTTGA.

### Eggshell and permeability barrier removal

For eggshell removal, embryos were collected from dissected adult *gsr-1 (m+,z-)* worms in egg buffer(118 mM NaCl, 48 mM KCl, 2 mM CaCl2, 2 mM MgCl2, 25 μM HEPES(pH7.4)). Developmental stages of collected embryos were not sorted out. Embryos of 2-cell-stage and later-stages were included. The embryos were incubated in 1% NaOCl for 2 min followed by the incubation in chitinase-chymotrypsine solution (2U/ml chitinase in egg buffer pH6.0) for 4 min at 22°C. It is noted that the permeability barrier inside the eggshell still remains around the embryos after this procedure.

For lethality measurements, the eggshell-removed embryos as well as intact *gsr-1 (m-, z-)* embryos were incubated in 100 μl of egg buffer in a chamber slide. After overnight incubation at 22 °C, unhatched embryos were counted under a stereomicroscope (Leica, M205C). DIC imaging was performed on an optical microscope (Olympus, BX51).

For the permeability barrier integrity assay, embryos were placed into a solution of 5 μg/ml FM4-64 in egg buffer and were mounted on a microscope slide. Fluorescent images were obtained with a confocal microscope (Nikon Ti-E with the spinning disk confocal unit CSU-X1(Yokogawa electric Corp.)) with a solid-state laser line (561 nm, Andor Technology).

### Graphical and Statistical Analysis

Data were processed in Excel (Microsoft Corporation) then Prism (GraphPad Software) was used to generate bar charts and perform the statistical analyses described in the Figure Legends.

## Data availability statement

*C. elegans* strains, *E. coli* strains and plasmids are available upon request. The authors affirm that all data necessary for confirming the conclusions of the article are present within the article, figures and tables with the exception of Supplementary Table 1 which contains *C. elegans* strains and genotypes used in this study. WGS data will be deposited at the Gene Expression Omnibus repository of NCBI (pending).

## Acknowledgements and funding

Some *C. elegans* strains were provided by the CGC, which is funded by NIH Office of Research Infrastructure Programs (P40 OD010440) USA and by the National BioResearch Project, Japan. We thank SunyBiotech (https://www.sunybiotech.com/) for their excellent assistance generating CRISPR-Cas9 edited alleles and the Genomics and Bioinformatic facility at CIBIR for their support. We also thank Profs Jan Gruber, Paul Sternberg, Gary Ruvkun and Chris Link for kindly providing strains.

M.V.V., D.G.G, E.G.O, J.C and A.M.V. were supported by Projects PGC2018-094276-B-I00, PID2021-122311NB-I00 and PID2021-127388NB-I00, financed by MICIU/AEI/10.13039/501100011033 and from FEDER, UE as well as by Projects P20_00229 and DOC_01674 Ayudas a proyectos I+D+I (PAIDI 2020) CTEICU – Junta de Andalucía. Co-financed with FEDER at 80%. Andalucía se mueve con Europa.

## Author contribution

A.M.V. and J.C. designed and supervised the research. M.V.V. and D.G.G. generated strains, performed *C. elegans* experiments and analyzed the data. E.G.O. performed qPCR analysis and analyzed the data. A.Z. performed single-nucleotide polymorphism (WGS-SNP) mapping strategy. Q.C. and E.S.J.A. performed all experiments with recombinant TRXR-1. J.A.M.L. identified *gnaIs2* transgene as suppressor of *gsr-1* mutants embryonic lethality. D.P. M.F. and J.C. contributed with reagents and strains. N.J.O. identified the microdeletion at *smg-3* locus. A.H. and S.O. performed the embryo eggshell and permeability barrier experiments. A.M.V. wrote the manuscript and all the authors edited and reviewed it.

