## Supplemental Figures for "The transsulfuration pathway suppresses the embryonic lethal phenotype of glutathione reductase mutants in *Caenorhabditis elegans*"

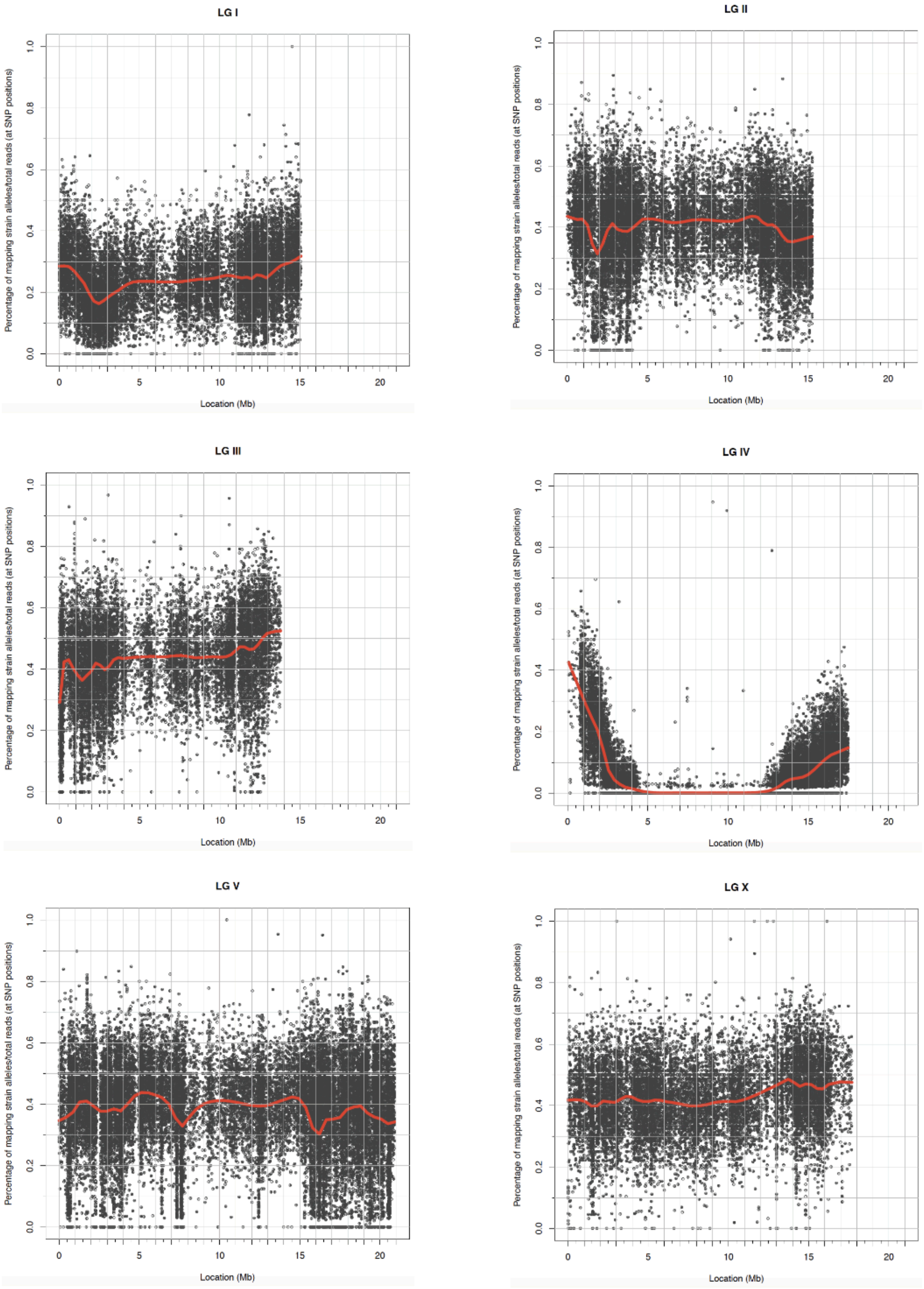

**Supplemental Figure 1. Identification of the chromosome harbouring the *gnals2* transgene.** CloudMap Hawaiian Variant Mapping with WGS tool plots the *gnals2* transgene in a large region of chromosome IV. The tool plots the ratio of *gnals2* alleles/total reads at each of the mapping gnals2 SNP positions in the genome. In red local regression line (LOESS) is plotted. LOESS regression is a locally weighted polynomial regression that gives weight to points near the position where *gnals2* is being estimated as well as to points further away on the chromosome.

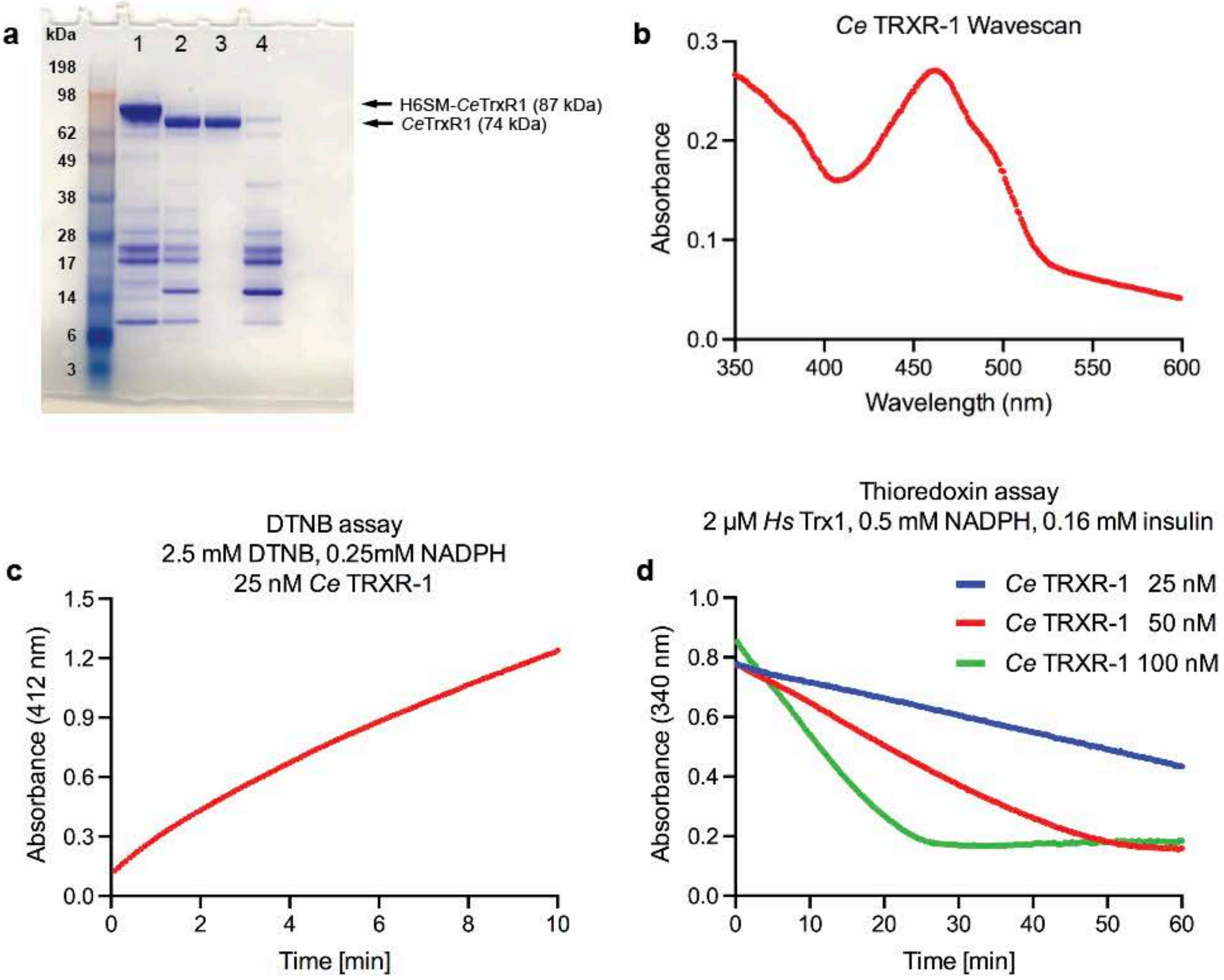

**Supplemental Figure 2. Biochemical characterization of recombinant *C. elegans* TRXR-1. a) SDS gel of *Ce*TRXR-1 protein fractions.** Lane 1) Purification of *Ce*TRXR-1 fused with N-terminal His-tag and SUMO tag (H6SM-*Ce*TrxR1) over a Nickel column (1st IMAC). Lane 2) Treatment of purified *Ce*TrxR1 fraction with SUMO protease ULP1, to remove the N-terminal tag. Lane 3) Flowthrough of the ULP1 treated sample loaded onto a Nickel column again (2nd IMAC) containing pure non-tagged *Ce*TrxR1. Lane 4) Elution of the Nickel column with imidazole, which contained the separated N-terminal H6SM-tag, ULP1 and impurities from the first IMAC. **b) UV-Visible wavescan spectrometry analysis of purified *Ce*TRXR-1.** The wavescan of purified *Ce*TRXR-1 shows a shoulder of absorbance around 450 nm, typical of proteins containing a FAD prosthetic group. **c) Determination of *Ce*TRXR-1 specific activity.** The DTNB reduction assay was used to determine the specific activity of purified *Ce*TRXR-1, which resulted in 2 Units/mg. **d) Thioredoxin reduction activity assay.** The *Ce*TRXR-1 preparation was assayed with increasing concentrations of human TRX1 and the consumption of NADPH was measured, demonstrating that the *Ce*TRXR-1 preparation is active as thioredoxin reductase.

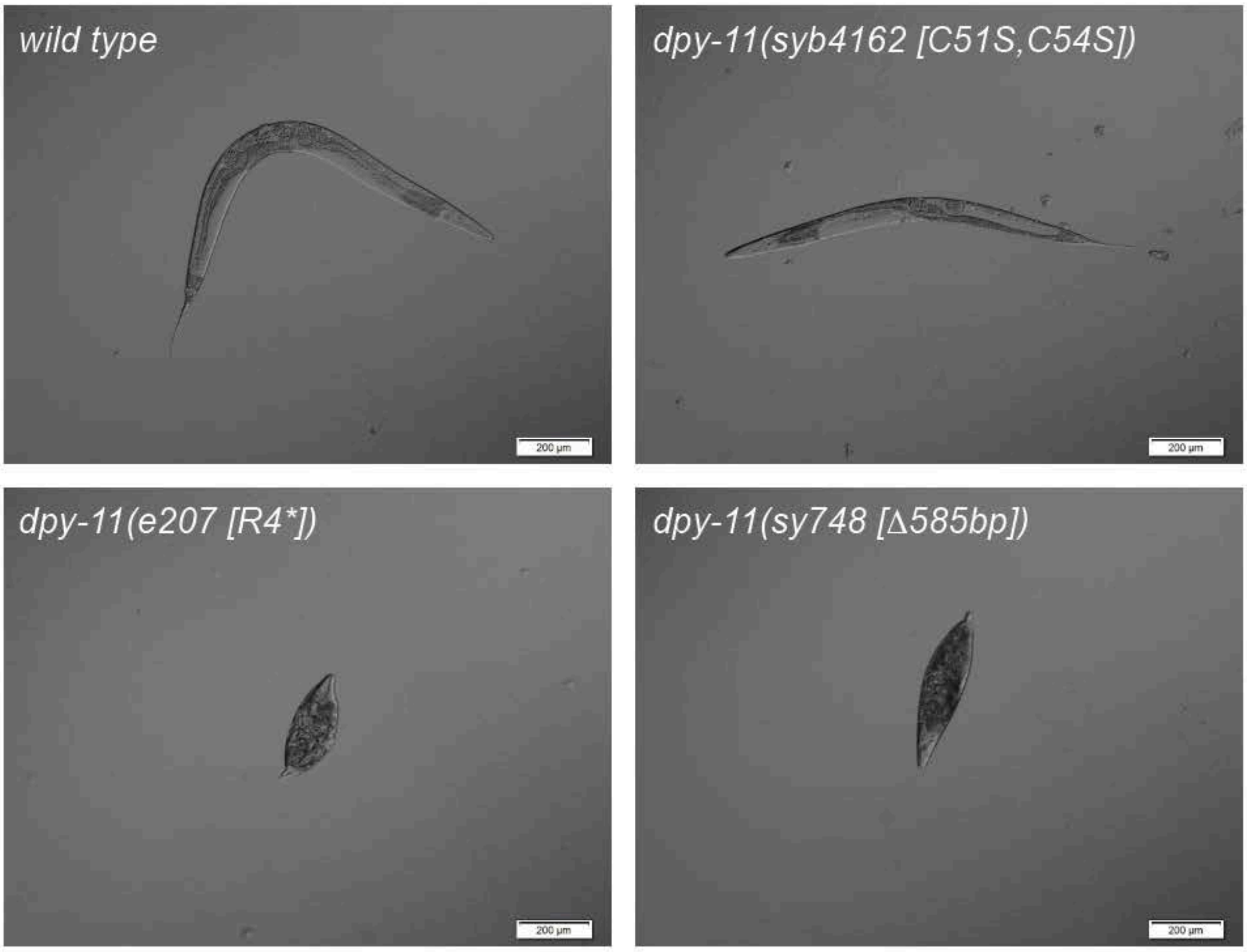

**Supplemental Figure 3. Representative micrographs of the different *dpy-11* mutant alleles and its comparison with *wild type* worms.** Differential interface contrast images were adquired at the first day adult stage. The asterisk in the *dpy-11(e207)* allele indicates a stop codon.

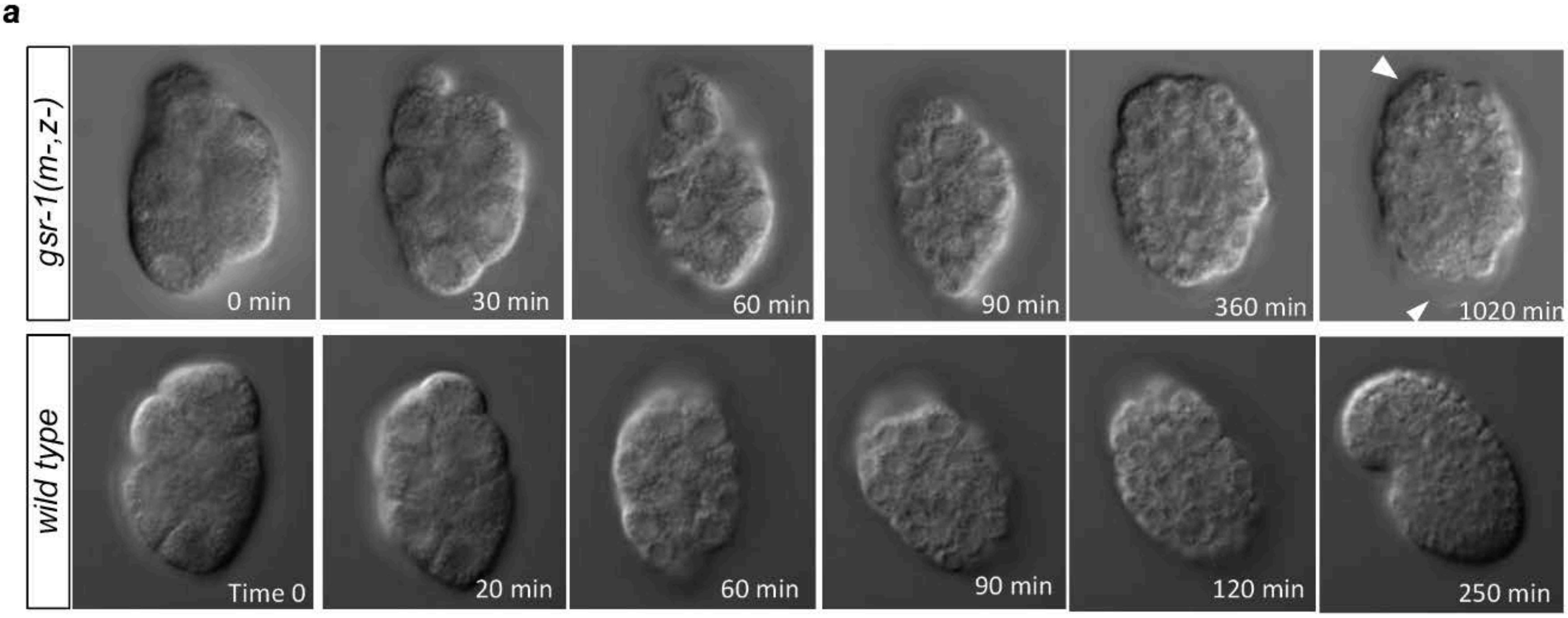

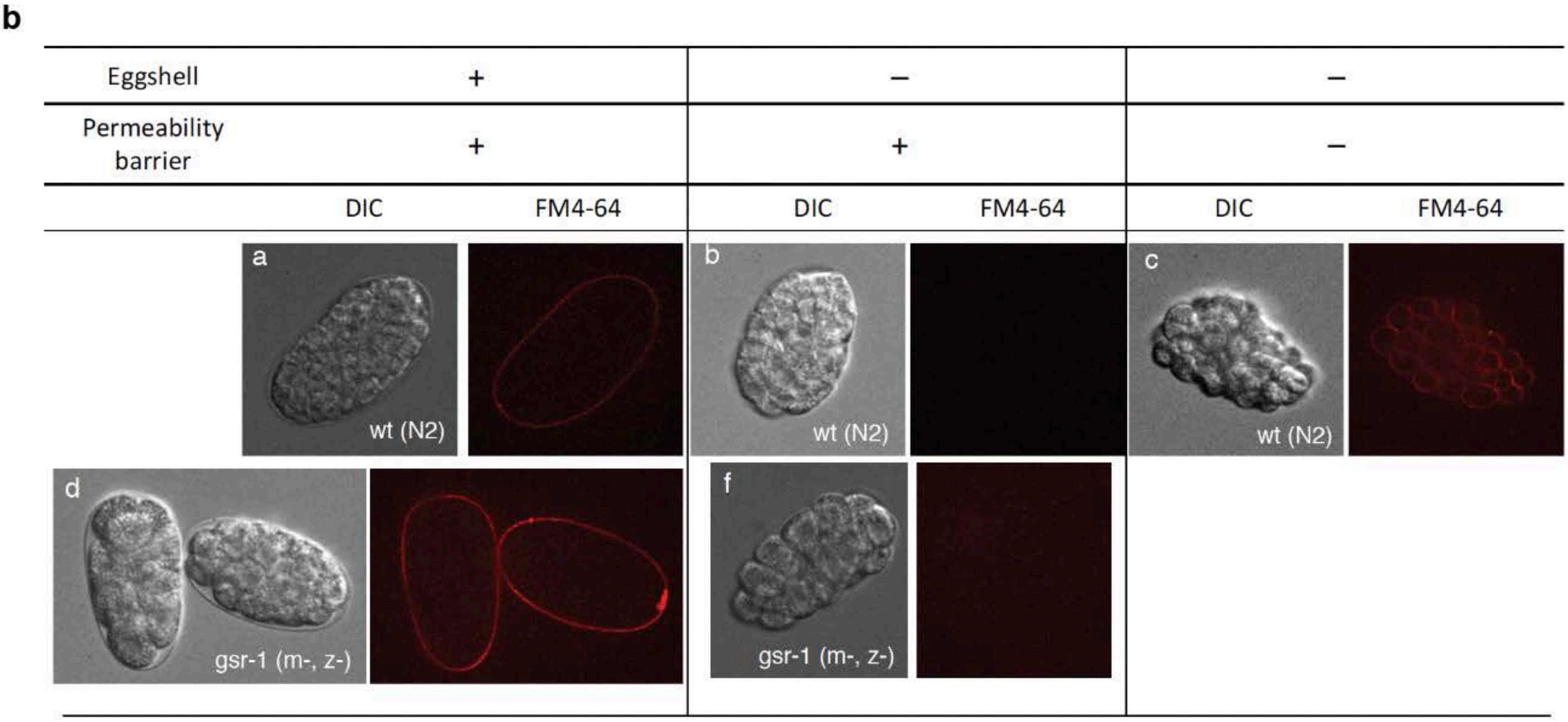

**Supplemental Figure 4. a) Differential interference contrast images of wild type and *gsr-1(m-,z-)* embryos after chitinase treatment.** Imaging started (time 0) 10 minutes after chitinase treatment. Deformed permeability barrier is observed in the *gsr-1* embryos (arrowheads) but not in *wild type* embryos.
**b) Dye assay to monitor the integrity of embryo permeability barrier.** *Wild type* and *gsr-1* embryos were exposed to FM4-64 dye. Embryos with intact eggshell do not incorporate the dye which accumulated surrounding the embryo externally. When the eggshell is removed, no staining is observed. Only when the permeability barrier is removed, the embryonic cells are stained. However, this treatment cause the arrest of embryonic development.

**Supplemental Table 1: *C. elegans* strains used in this study**

| Strain name | Genotype | Strain origin/construction |
|---|---|---|
| N2 | *Wild type, DR subclone of CB original (Tc1 pattern I)* | [a]CGC |
| VZ454 | *gsr-1(tm3574) / qC1 dpy-19(e1259) glp-1(q339) nIs281[Pmyo-2::RFP] III* | Mora-Lorca *et al.* (2016) Free Radic. Biol. Med. 96: 446-461 |
| GRU102 | *gnaIs2 [Pmyo-2::yfp; Punc-119::Abeta1-42]* | Fong *et al.* (2016) Sci. Rep. 6: 33781 |
| VZ730 | *gsr-1(tm3574) III; gnaIs2 [Pmyo-2::yfp; Punc-119::Abeta1-42]* | This study, GRU102 x VZ454 |
| GRU101 | *gnaIs1 [Pmyo-2::yfp]* | Fong *et al.* (2016) Sci. Rep. 6: 33781 |
| VZ712 | *gsr-1(tm3574) / qC1 dpy-19(e1259) glp-1(q339) nIs281[Pmyo-2::RFP] III; gnaIs1 [Pmyo-2::yfp]* | This study, GRU101 x VZ454 |
| CL2355 | *smg-1 (cc546) dvIs50 [pCL45 (snb-1::Abeta 1-42::3' UTR(long) + mtl-2::GFP] I* | Wu *et al.* (2006) J. Neurosci. 26: 13102-13113 |
| VZ567 | *smg-1 (cc546) dvIs50 [pCL45 (snb-1::Abeta 1-42::3' UTR(long) + mtl-2::GFP] I; gsr-1(tm3574) / qC1 dpy-19(e1259) glp-1(q339) nIs281[Pmyo-2::RFP] III* | This study, CL2355 x VZ454 |
| CB4354 | *smg-3(ma117) IV; him-5(e1490) V* | [a]CGC |
| VZ919 | *gsr-1(tm3574) III; smg-3(ma117) IV* | This study, CB4354 x VZ454 |
| FX5719 | *smg-3(tm5719) IV* | [b]NRBP |
| VZ920 | *gsr-1(tm3574) III; smg-3(tm5719) IV* | This study, FX5719 x VZ454 |
| FX5906 | *smg-3(tm5906) IV* | [b]NRBP |
| VZ921 | *gsr-1(tm3574) III; smg-3(tm5906) IV* | This study, FX5906 x VZ454 |
| CB3214 | *smg-1(e1228) him-2(e1065) I* | [a]CGC |
| VZ938 | *smg-1(e1228) him-2(e1065) I; gsr-1(tm3574) III* | This study, CB3214 x VZ454 |
| CB4043 | *smg-2(e2008) I; him-5(e1490) V* | [a]CGC |
| VZ939 | *smg-2(e2008) I; gsr-1(tm3574) III* | This study, CB4043 x VZ454 |
| CB4554 | *smg-1(e1228) smg-2(e2008) I; him-5(e1490) V* | [a]CGC |
| VZ922 | *smg-1(e1228) smg-2(e2008) I; gsr-1(tm3574) III* | This study, CB4554 x VZ454 |
| PHX2363 | *gsr-1(syb2363) / qC1 dpy-19(e1259) glp-1(q339) III* | This study, CRISPR-Cas9 editing on N2 |
| VZ930 | *gsr-1(syb2363) / qC1 dpy-19(e1259) glp-1(q339) nIs281[Pmyo-2::RFP] III* | This study, PHX2363 x VZ454 |
| VZ940 | *smg-1(e1228) him-2(e1065) I; gsr-1(syb2363) III* | This study, CB3214 x VZ930 |
| VZ941 | *smg-2(e2008) I; gsr-1(syb2363) III* | This study, CB4043 x VZ930 |
| VZ942 | *smg-3(ma117) IV; gsr-1(syb2363) III* | This study, CB4354 x VZ930 |
| VB1414 | *trxr-1(sv47) IV* | Stenvall *et al.* (2011) Proc. Natl. Acad. Sci. USA 108:1064-1069 |
| VZ492 | *gsr-1(tm3574) / qC1 dpy-19(e1259) glp-1(q339) nIs281[Pmyo-2::RFP] III; trxr-1 (sv47) IV* | This study, VB1414 x VZ454 |
| VZ496 | *smg-2(e2008) I; gsr-1(tm3574) / qC1 dpy-19(e1259) glp-1(q339) nIs281[Pmyo-2::RFP] III; trxr-1(sv47) IV* | This study, CB4043 x VZ492 |
| CER346 | *trxr-1(cer34 [Sec666Cys]) IV* | This study, CRISPR-Cas9 editing on N2 |
| VZ936 | *gsr-1(tm3574) / qC1 dpy-19(e1259) glp-1(q339) nIs281[Pmyo-2::RFP] III; trxr-1(cer34 [Sec666Cys]) IV* | This study, CER346 x VZ454 |
| VZ497 | *smg-2(e2008) I; gsr-1(tm3574) / qC1 dpy-19(e1259) glp-1(q339) nIs281[Pmyo-2::RFP] III; trxr-1(cer34 [Sec666Cys]) IV* | This study, CB4043 x VZ936 |
| CER374 | *trxr-1(cer55 [Sec666STOP]) IV* | This study, CRISPR-Cas9 editing on N2 |
| VZ937 | *gsr-1(tm3574) / qC1 dpy-19(e1259) glp-1(q339) nIs281[Pmyo-2::RFP] III; trxr-1(cer55 [Sec666STOP]) IV* | This study, CER374 x VZ454 |
| VZ498 | *smg-2(e2008) I; gsr-1(tm3574) / qC1 dpy-19(e1259) glp-1(q339) nIs281[Pmyo-2::RFP] III; trxr-1(cer55 [Sec666STOP]) IV* | This study, CB4043 x VZ937 |
| CB207 | *dpy-11(e207) V* | [a]CGC |
| VZ983 | *gsr-1(tm3574) / qC1 dpy-19(e1259) glp-1(q339) nIs281[Pmyo-2::RFP] III; dpy-11(e207) V* | This study, CB207 x VZ454 |
| VZ993 | *gsr-1(tm3574) III; smg-3(tm5719) IV; dpy-11(e207) V* | This study, FX5719 x VZ983 |
| PS6624 | *dpy-11(sy748) V* | Chiu *et al.* (2013) Genetics 195: 1167-1171 |
| VZ1013 | *gsr-1(tm3574) / qC1 dpy-19(e1259) glp-1(q339) nIs281[Pmyo-2::RFP] III; dpy-11(sy748) V* | This study, PS6624 x VZ454 |
| VZ1008 | *gsr-1(tm3574) III; smg-3(tm5719) IV; dpy-11(sy748) V* | This study, FX5719 x VZ1013 |
| PHX4162 | *dpy-11(syb4162 [Cys51Ser; Cys54Ser]) V* | This study, CRISPR-Cas9 editing on N2 |
| VZ1019 | *dpy-11(syb4162 [Cys51Ser; Cys54Ser]) V* | This study, PHX4162 outcrosed 2x with N2 |
| VZ1031 | *gsr-1(tm3574) / qC1 dpy-19(e1259) glp-1(q339) nIs281[Pmyo-2::RFP] III; dpy-11(syb4162 [Cys51Ser; Cys54Ser]) V* | This study, VZ454 x VZ1019 |
| VZ1032 | *gsr-1(tm3574) / qC1 dpy-19(e1259) glp-1(q339) nIs281[Pmyo-2::RFP] III; smg-3(tm5719) IV; dpy-11(syb4162 [Cys51Ser; Cys54Ser]) V* | This study, FX5719 x VZ1031 |
| VZ1 | *trx-1(ok1449) II* | Miranda-Vizuete *et al.* (2006) FEBS Lett. 580: 484-490 |
| VZ1075 | *trx-1(ok1449) II; gsr-1(tm3574) III; smg-3(tm5719) IV* | This study, VZ1 x VZ920 |
| VZ1053 | *txdc-17(syb4767) III* | Martí-Andrés *et al.* (2024) EMBO J., in press. |
| VZ1066 | *gsr-1(tm3574) txdc-17(syb4767) / qC1 dpy-19(e1259) glp-1(q339) nIs281[Pmyo-2::RFP] III* | This study, VZ454 x VZ1053 |
| VZ1076 | *gsr-1(tm3574) txdc-17(syb4767) III; smg-3(tm5719) IV* | This study, FX5719 x VZ1066 |
| FX2462 | *txl-1(tm2462) I* | [b]NRBP |
| VZ108 | *txl-1(tm2462) I* | This study, FX2462 outcrosed 6x with N2 |
| VZ1083 | *txl-1(tm2462) I; gsr-1(tm3574) III; smg-3(tm5719) IV* | This study, VZ108 x VZ920 |
| MRF03 | *cth-1(ok3319) V* | Zivanovic *et al.* (2019) Cell Metab. 30: 1152-1170 |
| VZ1188 | *gsr-1(tm3574) / qC1 dpy-19(e1259) glp-1(q339) nIs281[Pmyo-2::RFP] III; smg-3(tm5719) IV; cth-1(ok3319) V* | This study, FX5719 x MRF03 x VZ454 |
| GR2257 | *cth-2(mg599) II* | Warnhoff and Ruvkun (2019) Nat. Chem. Biol. 15: 480-488 |
| VZ1189 | *cth-2(mg599) II; gsr-1(tm3574) / qC1 dpy-19(e1259) glp-1(q339) nIs281[Pmyo-2::RFP] III; smg-3(tm5719) IV* | This study, FX5719 x GR2257 x VZ454 |
| VZ1134 | *gsr-1(tm3574) txdc-17/trp-14(syb4767) / qC1 dpy-19(e1259) glp-1(q339) nIs281[Pmyo-2::RFP] III; smg-3(tm5719) IV; cth-1(ok3319) V* | This study, FX5719 x MRF03 x VZ1066 |
| VZ1104 | *cth-2(mg599) / mIn1[dpy-10(e128) mIs14(Pmyo-2::GFP)] II; gsr-1(tm3574) txdc-17/trp-14(syb4767) III; smg-3(tm5719) IV* | This study, FX5719 x VZ1066 x VZ1094 |
| VZ1206 | *cth-2(mg599) II; smg-3(tm5719) IV; cth-1(ok3319) V* | This study, FX5719 x MRF03 x GR2257 |
| VZ1183 | *cth-2(mg599) II; gsr-1(tm3574) / qC1 dpy-19(e1259) glp-1(q339) nIs281[Pmyo-2::RFP] III; smg-3(tm5719) IV; cth-1(ok3319) V* | This study, VZ454 x VZ1206 |
| VZ1121 | *cth-2(mg599) / mIn1[dpy-10(e128) mIs14(Pmyo-2::GFP)] II; gsr-1(tm3574) txdc-17/trp-14(syb4767) III; smg-3(tm5719) IV; cth-1(ok3319) V* | This study, MRF03 x VZ1104 |
| VZ1066 | *gsr-1(tm3574) txdc-17/trp-14(syb4767) / qC1 dpy-19(e1259) glp-1(q339) nIs281[Pmyo-2::RFP] III* | This study, VZ454 x VZ1053 |
| VZ1076 | *gsr-1(tm3574) txdc-17/trp-14(syb4767) III; smg-3(tm5719) IV* | This study, FX5719 x VZ1066 |
| VZ1094 | *cth-2(mg599) / mIn1[dpy-10(e128) mIs14(Pmyo-2::GFP)] II* | This study, VB2616 x GR2257 |
| VB2616 | *gcs-1(ok436) / mIn1[dpy-10(e128) mIs14(myo-2::GFP)] II* | Outcrossed 8x with N2. Simon Tuck gift |
| PHX7115 | *cth-1(syb7115 [cth-1::3xFLAG::eGFP] V* | This study, CRISPR-Cas9 editing on N2 |
| VZ1175 | *cth-1(syb7115 [cth-1::3xFLAG::eGFP] V* | This study, PHX7115 outcrosed 2x with N2 |
| VZ1195 | *gsr-1(tm3574) / qC1 dpy-19(e1259) glp-1(q339) nIs281[Pmyo-2::RFP] III; cth-1(syb7115 [cth-1::3xFLAG::eGFP] V* | This study, VZ454 x VZ1175 |
| VZ1196 | *smg-3(tm5719) IV; cth-1(syb7115 [cth-1::3xFLAG::eGFP] V* | This study, FX5719 x VZ1175 |
| VZ1197 | *gsr-1(tm3574) III; smg-3(tm5719) IV; cth-1(syb7115 [cth-1::3xFLAG::eGFP] V* | This study, VZ920 x VZ1175 |
| PHX7086 | *cth-2(syb7086 [cth-2::3xFLAG::eGFP] II* | This study, CRISPR-Cas9 editing on N2 |
| VZ1172 | *cth-2(syb7086 [cth-2::3xFLAG::eGFP] II* | This study, PHX7086 outcrosed 2x with N2 |
| VZ1192 | *cth-2(syb7086 [cth-2::3xFLAG::eGFP] II; gsr-1(tm3574) / qC1 dpy-19(e1259) glp-1(q339) nIs281[Pmyo-2::RFP] III* | This study, VZ454 x VZ1172 |
| VZ1193 | *cth-2(syb7086 [cth-2::3xFLAG::eGFP] II; smg-3(tm5719) IV* | This study, FX5719 x VZ1172 |
| VZ1194 | *cth-2(syb7086 [cth-2::3xFLAG::eGFP] II; gsr-1(tm3574) III; smg-3(tm5719) IV* | This study, VZ920 x VZ1172 |

[a] CGC: *Caenorhaditis* Genetics Center

[b] NBRP: National Bioresource Project
